# Triggering the expression of a silent gene cluster from genetically intractable bacteria results in scleric acid discovery

**DOI:** 10.1101/265645

**Authors:** Fabrizio Alberti, Daniel J. Leng, Ina Wilkening, Lijiang Song, Manuela Tosin, Christophe Corre

**Affiliations:** Warwick Integrative Synthetic Biology Centre and School of Life Sciences, University of Warwick, Coventry, CV4 7AL, UK; Department of Chemistry, University of Warwick, Coventry, CV4 7AL, UK

## Abstract

Herein we report a strategy for the rapid and rational characterisation of novel microbial natural products from silent gene clusters. A conserved set of five regulatory genes was used as a query to search genomic databases and identify atypical biosynthetic gene clusters (BGCs). A 20-kb BGC from the genetically intractable *Streptomyces sclerotialus* bacterial strain was captured using yeast-based homologous recombination and introduced into validated heterologous hosts. CRISPR/Cas9-mediated genome editing was then employed to rationally inactivate the key transcriptional repressor and trigger production of an unprecedented class of hybrid natural products exemplified by (2-(benzoyloxy)acetyl)-*L*-proline, named scleric acid. Subsequent rounds of CRISPR/Cas9-mediated gene deletions afforded a selection of biosynthetic gene mutant strains which led to a plausible biosynthetic pathway for scleric acid assembly. Scleric acid and a key biosynthetic intermediate were also synthesised and used as authentic standards. The assembly of scleric acid involves two unique enzymatic condensation reactions that respectively link a proline and a benzoyl residue to each end of a rare hydroxyethyl-ACP intermediate. Scleric acid was then shown to exhibit moderate activity against *Mycobacterium tuberculosis*, as well as modest inhibition of the cancer-associated metabolic enzyme Nicotinamide N-methyltransferase (NNMT).

---

While actinomycete bacteria have been the foremost producers of antibiotics since the mid-1940s, in the last decade high-throughput DNA sequencing technologies and novel bioinformatics tools have highlighted an immense number of uncharacterised biosynthetic gene clusters (BGCs) predicted to direct the assembly of bioactive natural products.^1^ The presence of BGCs was not only revealed in actinomycete genomes but also in those of human commensal and pathogenic bacteria (i.e. *Staphylococcus lugdinensis, Burkholderia cepacia* complex) as well as in the genomes of unculturable bacteria and in diverse metagenomic libraries.^2-6^

Despite the conspicuous number of specialised metabolites isolated from actinomycetes, only a small fraction of the natural products ‘encrypted’ at the DNA level has been exploited to date. Experimental characterisation of the biosynthetic product of a BGC is often laborious and time-consuming particularly due to the uniqueness of every microorganism. Protocols for introducing DNA into bacterial cells are species-dependent and often ineffective. Their optimisation can take years but many culturable micro-organisms remain genetically intractable. In any case the vast majority of bacteria is genetically intractable as they are still unculturable in the laboratory environment thus preventing the exploitation of BGCs using many of the previously reported strategies.^1,7^

In addition, the biosynthesis of specialised metabolites is often tightly controlled at the transcriptional level. Cluster-associated transcriptional regulators that belong to the TetR-family of transcriptional repressors are particularly numerous.^8^ Deletions of cluster-specific TetR-like transcriptional repressors have been shown to trigger overproduction of the corresponding specialised metabolites, as seen for the antibiotics methylenomycin and coelimycin in *Streptomyces coelicolor* A3(2) and for the urea-containing gaburedins in *Streptomyces venezuelae*^9-12^.

Genetic manipulations of *Streptomyces* genomes has classically been accomplished using established but often laborious protocols optimised for a specific bacterial strains.^13^ In recent years however, targeted genome editing has been revolutionised by the advent of clustered regularly interspaced short palindromic repeats (CRISPR)/CRISPR-associated (Cas) systems, which allow for generating clean genomic deletions/insertions.^14^ Toolkits for editing streptomycete genomes have been developed,^15-17^ enabling researchers to overcome the issues associated with classical methods of gene disruption, in particular when multiple mutation events are desirable. The number of available selectable markers and issues with potential restoration of the wild-type configuration due to occurrence of single-crossover events were notable limitations.

Here, we report a strategy for selecting atypical BGCs and for rapid and rational characterisation of the corresponding natural products. The BGC is first captured and transferred into a validated heterologous host where CRISPR/Cas9-mediated genome editing can be employed to efficiently de-repress the expression of silent BGCs (Fig. 1). We applied this approach for the identification, isolation and structural elucidation of a novel structural class of natural products from a silent and cryptic gene cluster found in the soil-dwelling species *Streptomyces sclerotialus* NRRL ISP-5269, a species of filamentous bacteria first isolated in Poona (India).^18^ Beyond the discovery of this specialised metabolite, we strongly believe our approach will accelerate future research aimed at the characterisation of cryptic gene clusters from genetically intractable bacterial species or even from metagenomes.

**Figure 1.**
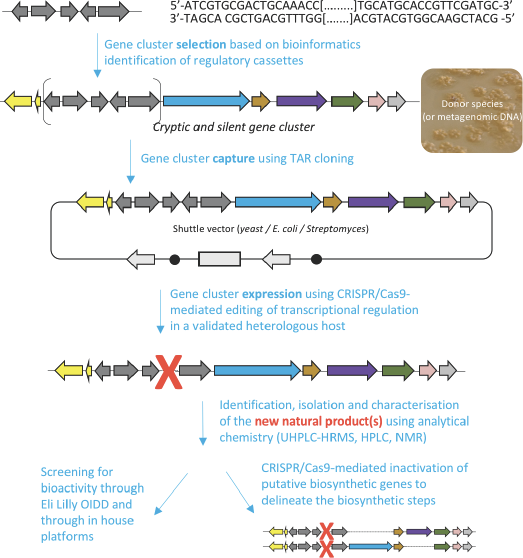
Overview of the workflow used in this study to characterise novel natural products from cryptic and silent gene clusters.

## Results

### Identification of the *scl* gene cluster in *S. sclerotialus* NRRL ISP-5269

The prioritisation of the gene cluster under study was guided by the presence of a specific set of five regulatory genes that we had previously characterised and exploited to trigger the expression of a silent and cryptic biosynthetic gene cluster in *Streptomyces venezuelae*.^12^ A Multigene BLAST^19^ search was undertaken using as a query the DNA sequences of the following five genes: *mmyR, mmfR* (both code for TetR-like transcriptional repressors) and *mmfLHP,* responsible for the biosynthesis of signalling molecules, known as methylenomycin furans (MMF) that trigger production of the methylenomycin antibiotics in *S. coelicolor* A3(2).^9^ In previous studies we have shown that inactivation of the *mmyR*-like transcriptional repressors is an effective approach for de-repressing silent gene clusters in actinomycetes.^10-12^

Consequently, orthologues of the methylenomycin transcriptional regulators and their associated regulatory proteins were identified. Among the hits generated, the gene cluster from *S. sclerotialus* NRRL ISP-5269, named hereafter *scl* cluster, was chosen for further study; the nucleotide sequence containing the *scl* cluster was available from the GenBank accession number JOBC01000043.1. In addition to homologues of the five genes used as query, the genetic organisation of the *scl* gene cluster included two adjacent and divergent operons of biosynthetic genes (Fig. 2a). A combination of AntiSMASH^20^ and manual BLASTp^19^ analyses indicated the putative borders of the *scl* cluster (Fig. 2a and Table 1). The predictive power of these modern bioinformatics tools often permits to deduce the chemical structure(s) of cryptic gene cluster product, particularly when modular systems such as type I modular polyketide synthases (PKS) or non-ribosomal peptide synthetases (NRPS) direct the biosynthesis.^22^ However, the originality of the *scl* cluster prevented such predictions and we worked on the assumption that a lack of bioinformatics prediction was more likely to result in a more structurally diverse and therefore truly novel natural product. The cluster spanned a region of 19,782 bp, and comprised 18 putative genes: 11 biosynthetic genes, 6 genes for regulation and 1 gene coding for a membrane transporter (Fig. 2a and Table 1).

**Figure 2.**
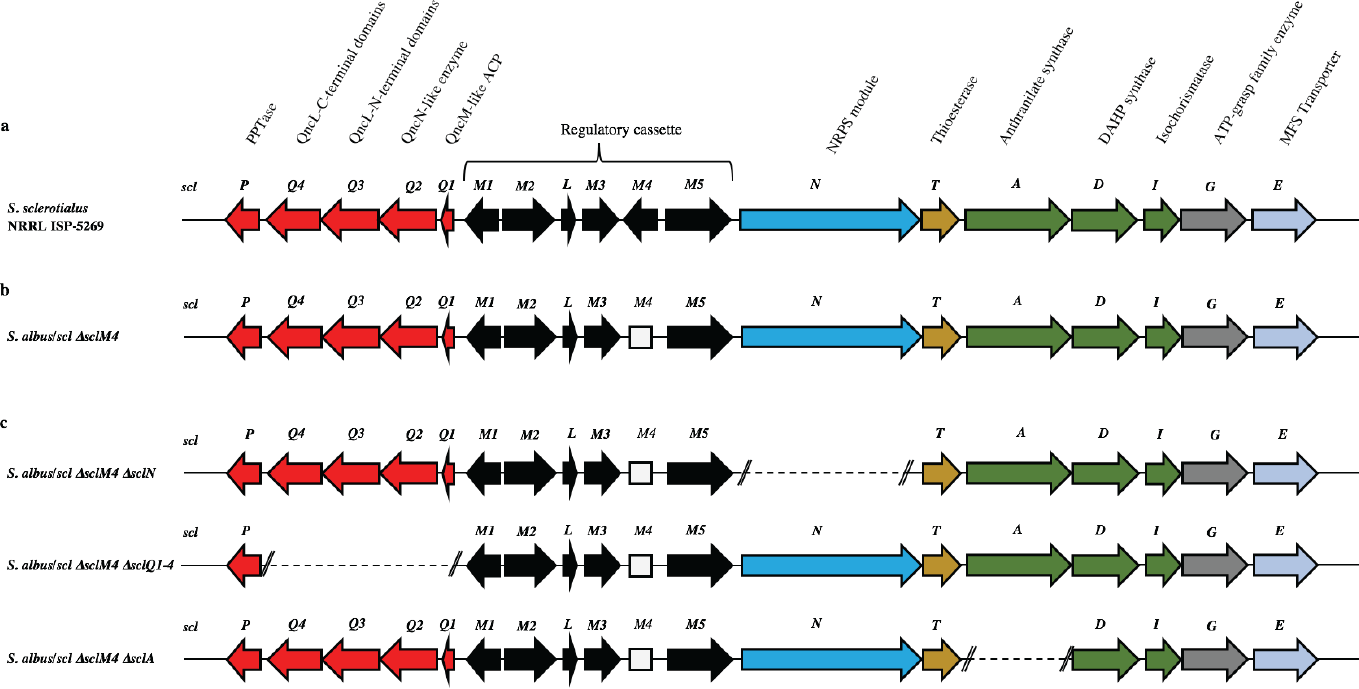
Genetic organisation of the *scl* gene cluster from *S. sclerotialus* NRRL ISP-5269 characterised in this study and schematic representation of mutants generated. (**a**) Gene cluster (19,782 bp) sequenced from *S. sclerotialus* NRRL ISP-5269. (**b**) Configuration of the *scl* gene cluster after CRISPR/Cas9-mediated targeted gene deletion within the heterologous host *S. albus.* The 20-bp out of frame deletion of *sclM4* generated a truncated gene, represented here as a rectangular shape for the derived gene sequence. (**c**) Configuration of the *scl* cluster within *S. albus* double mutants; deletion of genes *sclN, sclQ1-4* and *sclA* were generated starting from strain *S. albus/scl ΔsclM4,* and are represented here as dotted lines.

**Table 1.**
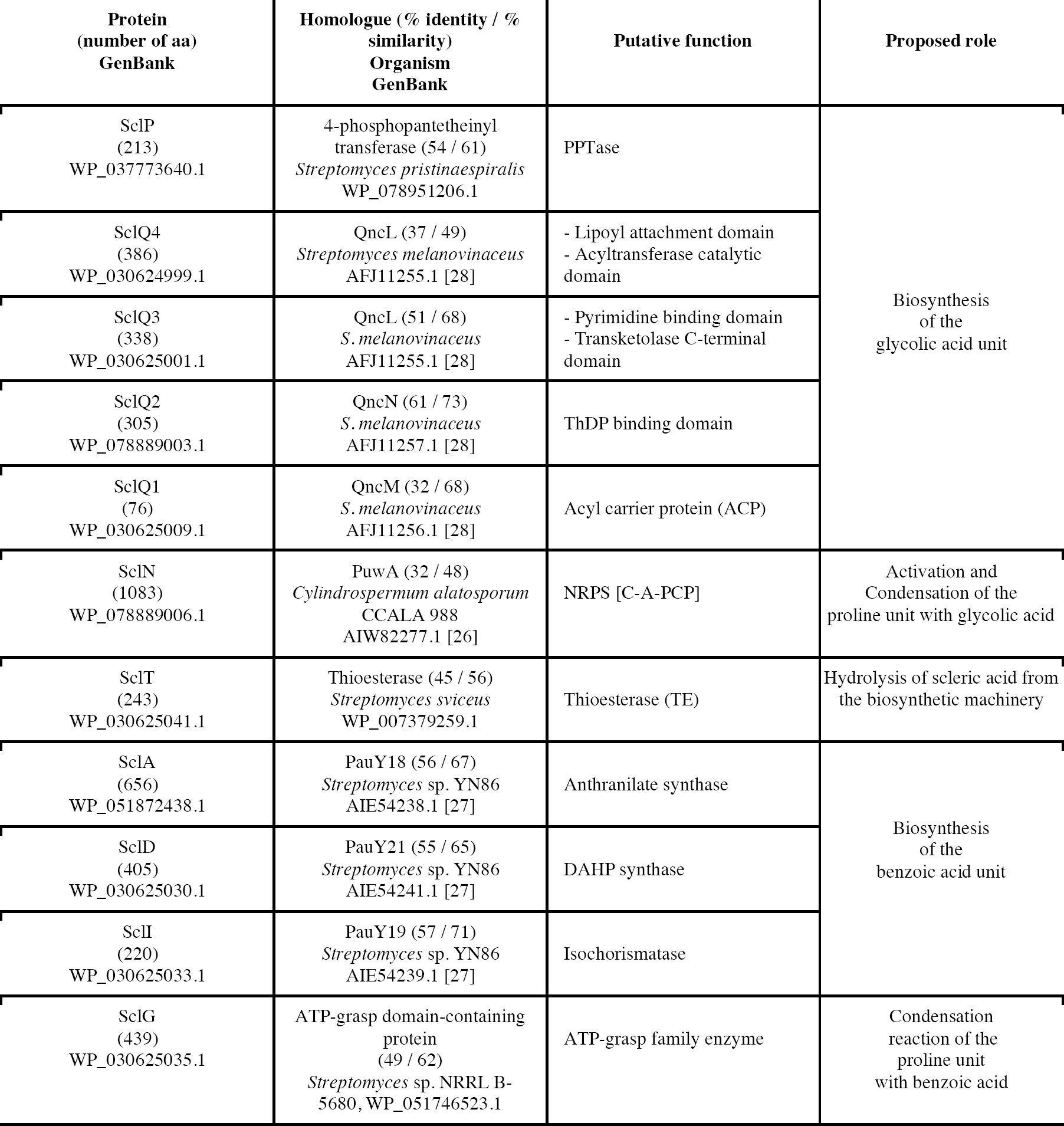
Proposed function of the biosynthetic genes from the *scl* cluster.

### Capture of the *scl* gene cluster and introduction into heterologous hosts

Introduction of plasmid DNA into *S. sclerotialus* NRRL ISP-5269 was attempted *via* intergenic-conjugation of mycelia with *E. coli* ET12567/pUZ8002 or protoplast transformation but these processes were found ineffective.^13^ The preparation of *S. sclerotialus* NRRL ISP-5269 spores was also attempted but despite screening various culture media in the aim to induce sporulation, this strain showed little aerial growth. We therefore set out to capture and heterologously express the *scl* gene cluster, with the aim of characterising its biosynthetic product(s) in a more genetically amenable host. A 33-kb region of genomic DNA that included the 19.7-kb *scl* BGC was captured from the purified genomic DNA of *S. sclerotialus* ISP-5269 *via* transformation-associated recombination (TAR) cloning.^23^ A pCAP03-derived plasmid was first assembled to specifically recombine in yeast with both extremities of the 33-kb target DNA fragment.^24^ The pCAP03-*scl* construct was introduced and stably integrated into the genome of *Streptomyces albus* J1074 and *S. coelicolor* M1152 *via* intergenic tri-parental conjugation with *E. coli* ET12567/pCAP03-scl and *E. coli* ET12567/pUB307. In parallel, the negative control strains *S. albus*/pCAP03 and *S. coelicolor* M1152 /pCAP03 were generated using the “empty” pCAP03 plasmid. The four *Streptomyces* strains were grown on supplemented minimal agar medium for 5 days, and the acidified agar medium was extracted with ethyl acetate. Their metabolic profiles were analysed by ultra-high pressure liquid chromatography-high resolution mass spectrometry (UHPLC-HRMS). Comparison of the MS chromatograms failed to reveal any new compounds in the heterologous hosts where the *scl* BGC had been integrated.

### De-repression of the *scl* gene cluster and characterisation of scleric acid

The lack of accumulation of novel metabolites in the *scl*-containing strains was likely to be due to the transcriptional repression activity of the TetR-like regulators encoded in the *scl* cluster. We had previously shown that genetic inactivation of *mmyR-*like genes in *S. coelicolor* A3(2) and in *S. venezuelae* would specifically de-repress the expression of adjacent BGCs.^10,12^ We therefore decided to genetically inactivate the *mmyR* homologous gene, *sclM4* using the CRISPR/Cas9-based plasmid pCRISPomyces2 (pCm2).^15^ For this purpose, we assembled the plasmid pCm2-*sclM4* and attempted intergenic-conjugation of *S. albus/scl* and *S. coelicolor* M1152/scl with *E. coli* ET12567/pUZ8007/pCm2-sclM4. Ex-conjugants for strain *S. albus/scl ΔsclM4* (see Fig. 2b for a representation of the genotype) were readily obtained, however no ex-conjugants could be obtained when attempting insertion of plasmid pCm2-sclM4 into *S. coelicolor* M1152/scl. The desired 20-bp out-of-frame deletion of *sclM4* was confirmed by sequencing of a PCR product using *S. albus/scl ΔsclM4* genome as a template (Supplementary Fig. S1). Metabolites produced by this strain were compared by UHPLC-HRMS to those of *S. albus/scl* and *S. albus/pCAP03* grown under the same conditions as those described earlier. *S. albus/scl ΔsclM4* showed accumulation of a major metabolite (Fig. 3a-c) with a retention time of 16.4 minutes on C_18_ reverse phase HPLC column and an *m/z* value of 278.1020 [M(C_14_H_15_NO_5_) + H]+ (calculated *m/z* of 278.1023). This compound was purified using a combination of flash chromatography on C_18_-silica column and HPLC. Its structure was then elucidated by a combination of 1D- and 2D-NMR spectroscopy experiments (Supplementary Figs. S2-6). The novel compound was characterised as being (2-(benzoyloxy)acetyl)-*L*-proline and named scleric acid (Fig. 3d).

**Figure 3.**
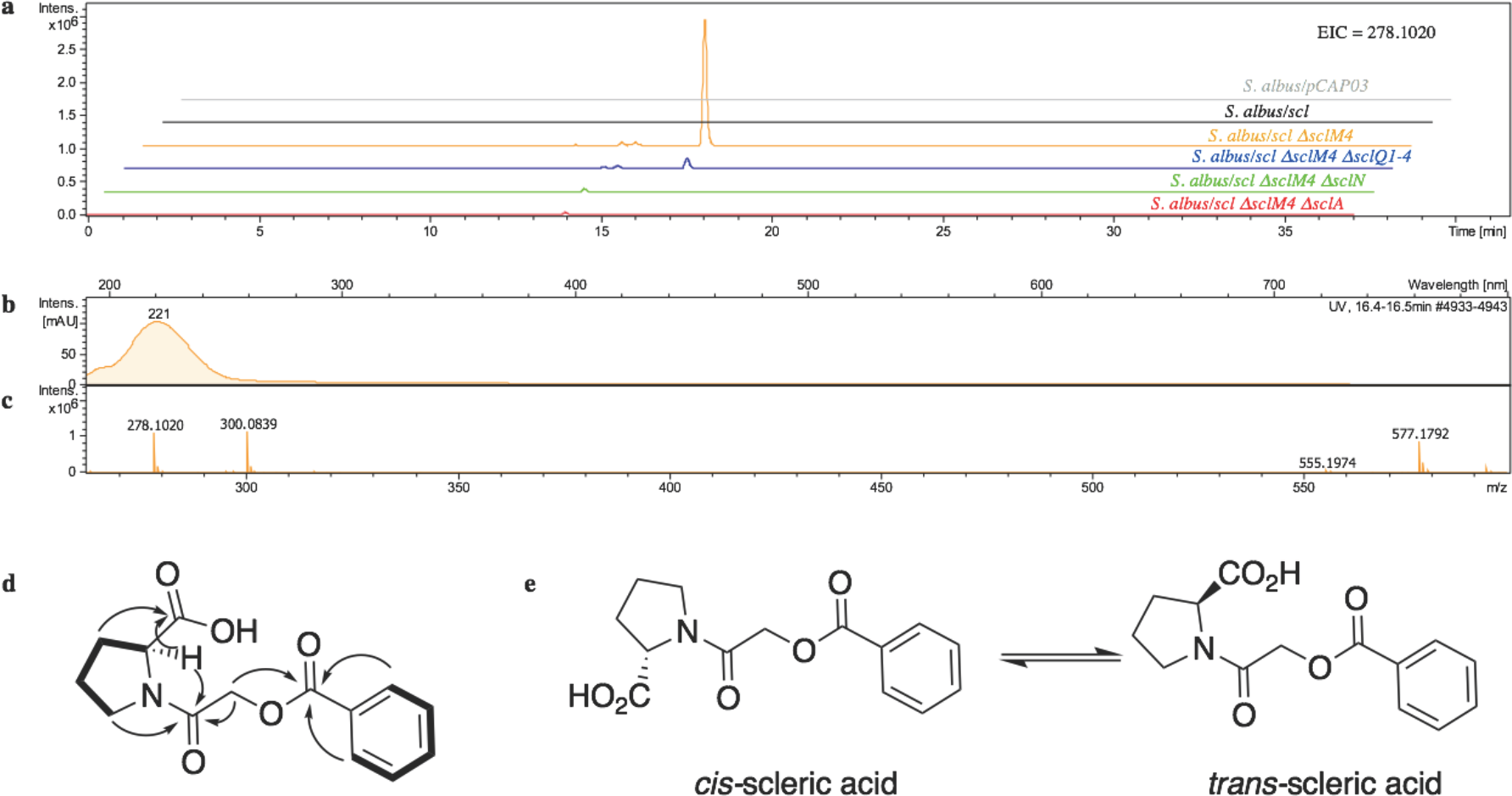
Identification and characterisation of scleric acid. (**a**) UHPLC-HRMS detection of metabolites produced in *S. albus/pCAP03* (grey trace), *S. albus/scl* (black), *S. albus/scl ΔsclM4* (orange), *S. albus/scl ΔsclM4 ΔsclQ1-4* (blue), *S. albus/scl ΔsclM4 ΔsclN* (green) and *S. albus/scl ΔsclM4 ΔsclA* (red). Extracted ion chromatograms in positive mode for *m/z* = 278.1020 are shown, highlighting accumulation of a new metabolite (scleric acid) at retention time 16.4 minutes in *S. albus/scl ΔsclM4* (orange trace). (**b**) UV chromatogram of scleric acid. (**c**) High-resolution mass spectrum in positive mode of scleric acid. (**d**) Selected correlations observed in the COSY (bold lines) and HMBC (arrows) NMR spectra of scleric acid. (**e**) Chemical structure of scleric acid that can adopt two main rotamer conformations.

In order to establish the stereochemistry of the proline residue, scleric acid was hydrolysed and derivatised with Marfey’s reagent.^25^ *L-* and D-proline were also derivatised using the same procedure and used as standard for HPLC comparison. Approximately 95% of the proline residue of scleric acid purified from *S. albus/scl ΔsclM4* was found to correspond to L-proline (Supplementary Fig. S7).

To confirm the proposed structure of scleric acid, an authentic standard was synthesised. A structural analogue, with the three building blocks rearranged and possibly consistent with the initial NMR data obtained, was also synthesised (Supplementary Figs. S8-12). LC-MS analyses and NMR data of both of these compounds unequivocally confirmed the proposed structure for scleric acid; the analogue revealed different NMR spectra and its physico-chemical properties resulted in a different retention time on LC-MS.

### Defining key biosynthetic genes in the *scl* cluster

BLASTp analyses^19^ combined with the elucidated structure for scleric acid allowed us to propose plausible functional assignments for all the biosynthetic enzymes coded in the *scl* BGC (Table 1).^26-28^

SclQ1-4 showed high homology to a set of three enzymes – QncN, QncL and QncM – from *Streptomyces melanovinaceus.* This set of enzymes has been shown to direct the biosynthesis and attachment of a C_2_-glycolicacyl unit to a NRP.^28^ More specifically, SclQ2 was homologous to QncN, a thiamin diphosphate (ThDP) binding domain. SclQ3 was homologous to the first two N-terminal domains of QncL, a pyruvate dehydrogenase/transketolase pyrimidine binding domain and a transketolase C-terminal domain while SclQ4 was homologous to the last two domains of QncL, a lipoyl attachment domain and an acyltransferase catalytic domain. Lastly, SclQ1 was homologous to the acyl carrier protein (ACP) QncM.

SclA showed high similarity to anthranilate synthase enzymes, and together with SclD (putative DAHP synthase) and SclI (putative isochorismatase) was hypothesised to be responsible for biosynthesis of the benzoyl group found in scleric acid *via* chorismate as an intermediate.

SclN was predicted to be a non-ribosomal peptide synthetase (NRPS) enzyme consisting of a single minimal elongation module: a putative, atypical condensation domain (C), an adenylation domain (A) and a peptidyl carrier protein (PCP) domain.^29^ The SclN A-domain was predicted to specifically activate L-proline, which was in accordance with the presence of a L-proline residue in scleric acid.^21^ The SclN C-domain was proposed to catalyse the amide bond formation between *L*-proline and glycolic acid.

The role of SclN, SclA, and SclQ1-4 in the biosynthesis of scleric acid was investigated by constructing gene deletion mutants in strains where the transcriptional repressor *sclM4* has also been inactivated *(S. albus/scl ΔsclM4* background). Plasmids pCm2-*sclN*, pCm2-*sclA* and pCm2-*sclQ1-4* were assembled and used to generate double mutant strains *S. albus/scl ΔsclM4 ΔsclN, S. albus/scl ΔsclM4 ΔsclA* and *S. albus/scl ΔsclM4 ΔsclQ1-4.* Deletions were confirmed by PCR screening (Supplementary Fig. S13). UHPLC-HRMS analysis revealed that production of scleric acid was abolished in *S. albus/scl ΔsclM4 ΔsclN* and *S. albus/scl ΔsclM4 ΔsclA* (Fig. 3a), confirming the essential role of SclN and SclA. Residual scleric acid production was detected from *S. albus/scl ΔsclM4 ΔsclQ1-4*; this could be explained by the fact that glycolic acid is known to be produced by *Streptomyces* species in particular for the biosynthesis of N-glycolylmuramic acid.^30^ Addition of 5 mM glycolic acid to the culture medium of *S. albus/scl ΔsclM4 ΔsclQ1-4* also resulted in scleric acid being produced in similar level to that observed with *S. albus/scl ΔsclM4* (Supplementary Fig. S14).

The identification of key precursors in scleric acid biosynthesis was also exploited to further increase the titres of scleric acid produced by *S. albus/scl ΔsclM4.* Enriching the culture medium with 5 mM *L*-proline, 5 mM benzoic acid or 5 mM glycolic acid significantly increased levels of scleric acid observed upon UHPLC-HRMS analysis of the ethyl acetate extracts compared to those observed with *S. albus/scl ΔsclM4* grown on the standard supplemented minimal medium (Supplementary Fig. S15).

Based on the predicted function of the *scl* biosynthetic genes, we therefore proposed a putative biosynthetic route to scleric acid as shown in Fig. 4 and described in the discussion.

**Figure 4.**
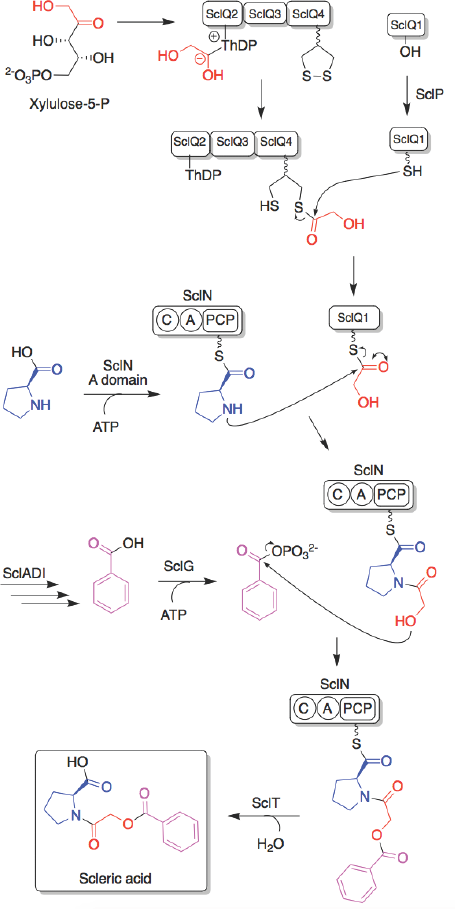
Proposed biosynthetic pathway to scleric acid.

### Biological activity of scleric acid

The antimicrobial activity of scleric acid was assessed. Inhibitory activity against representatives of the ESKAPE panel of pathogens: *Enterococcus faecium, Staphylococcus aureus, Klebsiella pneumoniae, Acinetobacter baumannii, Pseudomonas aeruginosa,* and *Enterobacter cloacae* was first screened but all strains appeared to be resistant to scleric acid, giving no observable MIC (Supplementary Table S1).

Scleric acid was then tested for a broader range of pharmaceutically relevant bioactivities through the Eli Lilly Open Innovation Drug Discovery (OIDD) Program. In a single point (20 μM) primary assays, scleric acid showed moderate antibacterial activity against *Mycobacterium tuberculosis* (H37Rv), exhibiting a 32% inhibition on the growth of this strain.

Scleric acid showed modest inhibitory activity on the cancer-associated metabolic enzyme Nicotinamide N-methyltransferase (NNMT), whose overexpression is known to contribute to tumorigenesis.^31^ NNMT catalyses the transfer of a methyl group from *S*-adenosyl-*L*-methionine (SAM) to nicotinamide, generating *S*-adenosyl-*L*-homocysteine (SAH) and 1- methylnicotinamide (MNAN).^31^ Scleric acid showed, on a concentration response curve assay, IC50 of 178.0 µM (NNMT MNAN) and 186.6 µM (NNMT SAH) (Supplementary Figure S16).

## Discussion

Specialised metabolites from actinomycete bacteria are one of the most valuable sources of novel antibiotics, as well as of other useful bioactive compounds employed in various fields, from human medicine to crop protection. High-throughput sequencing of bacterial genomes/metagenomes has become quick and inexpensive, and is unearthing a myriad of putative gene clusters that are awaiting to be characterised and exploited.

In this study, we developed a pipeline for rational characterisation of specialised metabolites from actinomycete genomes (Fig. 1). Starting with the bioinformatics identification and selection of cryptic gene clusters containing a characterised regulatory system, a DNA fragment containing the entire gene cluster is then captured *via* TAR cloning and introduced in the genome of a validated heterologous host. Expression of the biosynthetic genes are then triggered through CRISPR/Cas9-mediated editing of the key *mmyR-*like transcriptional repressors. Analytical chemistry procedures are then undertaken to identify, isolate and characterise the new natural product(s), overproduced in the heterologous host. Finally, subsequent rounds of CRISPR/Cas9-mediated gene deletions target putative biosynthetic genes and afford mutant strains. In turn, a biosynthetic route to the novel natural products can be proposed.

Our approach was validated by the discovery of (2-(benzoyloxy)acetyl)-*L*-proline, named scleric acid, from the genome of the soil-dwelling filamentous bacterium *Streptomyces sclerotialus* NRRL ISP-5269. The gene cluster of interest (*scl*) was identified *via* Multigene BLAST analysis, using as query the five-gene regulatory cassette involved in regulation of biosynthetic gene expression for methylenomycin production in *Streptomyces coelicolor* A3(2).^10^ The *scl* cluster was captured using TAR cloning and transferred into two different heterologous hosts: *Streptomyces albus* J1074 and *Streptomyces coelicolor* M1152. These two strains were chosen as they are well characterised chassis for heterologous expression of actinomycete gene clusters.^32,33^ While no novel metabolite was observed in either of the two hosts, CRISPR/Cas9-mediated deletion of the *sclM4* transcriptional repressor, homologous to the *tetR-like* repressor *mmyR* from the methylenomycin gene cluster,^10^ was attempted in the heterologous hosts. Mutant strains of *S. albus/scl* were readily obtained, whereas *S. coelicolor* M1152/scl failed to return any ex-conjugants using the same approach. The inefficacy of pCRISPomyces2 to edit the genome of *S. coelicolor* was unclear, however alternative CRISPR/Cas9-based methods for editing of this model species have now been described in the literature.^16-17^ De-repression of the *scl* cluster upon inactivation of the TetR-like repressor SclM4 within *S. albus/scl* resulted in accumulation of a novel hybrid compound, named scleric acid. Two sets of NMR signals were observed for the purified natural product as well as for the synthetic standard and revealed that scleric acid existed as two different rotamers, trans- and cis-scleric acid (Fig. 3e and Fig. S2). This is consistent with literature data for synthetic N-benzoyl-L-proline methyl ester where a 4:1 mixture of the two rotamers was observed.^34^

A second round of CRISPR/Cas9-based editing was undertaken on *S. albus/scl ΔsclM4* to inactivate putative key biosynthetic genes. This work established that the non-ribosomal peptide synthetase (NRPS) gene *sclN* and the anthranilate synthase-like gene *sclA* are essential for scleric acid assembly. Deletion of the four-gene operon *sclQ1-4* did not completely abolish scleric acid production, which could be explained by the fact that *Streptomyces* species are known to produce glycolic acid by other means.^30^

As expected enrichment of the culturing medium with the key biosynthetic precursors *i.e. L*-proline, benzoic acid and glycolic acid led *S. albus/scl ΔsclM4* to produce scleric acid in higher titres than when cultured in the supplemented minimal medium (Supplementary Fig. S15). The strategy of manipulating the pathway-specific transcriptional regulatory system also makes scleric acid production not reliant on a complex culture medium. Importantly the utilisation of supplemented minimal media does significantly facilitate the isolation of the natural product of interest.

Based on the predicted function of the Scl enzymes a plausible biosynthetic route to scleric acid was proposed (Fig. 4). A glycolic acid unit is believed to be produced by SclQ2-4 and loaded onto the SclQ1 ACP. SclP would convert apo-SclQ1 into holo-SclQ1 by phosphopantetheinylation. Meanwhile the adenylation domain of the NRPS SclN would activate *L*-proline, which could then be condensed to the glycolic acid unit through the activity of the C-domain of SclN. The three enzymes SclADI would then direct the biosynthesis of the benzoic acid moiety which would be activated by the ATP-grasp family enzyme SclG. That same enzyme would also catalyse the condensation of the benzoyl unit with *L*-proline-oxyacetyl-SclQ1. Lastly, scleric acid would be released as the product by the thioesterase SclT and exported out of the cell by the putative MFS transporter SclE.

In addition to the novel biochemistry this biosynthetic pathway offers, scleric acid has been shown to exhibit moderate antibacterial activity against *M. tuberculosis,* as well modest inhibition on the cancer-associated enzyme Nicotinamide N-methyltransferase (NNMT). We are currently investigating scleric acid bioactivity further through the Eli Lilly Open Innovation Drug Discovery (OIDD) Program.

In conclusion, the pipeline we developed represents a rational and rapid approach for the characterisation of novel microbial natural products and can be employed for the discovery of new metabolites from cryptic and silent gene clusters that originate from genetically-intractable or pathogenic bacterial species.

## Methods

### Bioinformatics analysis

The whole genome sequence of *Streptomyces sclerotialus* NRRL ISP-5269 was downloaded from the Genomes Online Database of the JGI Portal (U.S. Department of Energy, https://gold.jgi.doe.gov/, GOLD Project ID Gp0187859). Contig43 of the genome contained the *scl* gene cluster object of this study. A combination of AntiSMASH^20^ and manual BLASTp^19^ analyses allowed to define the putative borders of the *scl* cluster and predict the function of the encoded enzymes. The website for PKS/NRPS analysis from the University of Maryland was used for domain prediction of SclN.^22^ SerialCloner 2.6.1 (SerialBasics) was used for DNA sequence analysis and plasmid design.

### Reagents

All chemicals were purchased from Sigma-Aldrich, unless otherwise stated. Phusion DNA polymerase, as well as all restriction endonucleases, T4 DNA ligase, shrimp alkaline phosphatase (rSAP) and Gibson Assembly cloning kit, were purchased from New England Biolabs. Zymolyase 20T was purchased from MP Biomedicals, 5-fluoroorotic acid (5-FOA) was purchased from Thermo Fisher Scientific.

### Culturing and engineering of microorganisms

*Streptomyces sclerotialus* NRRL ISP-5269 was obtained from JCM (Japan Collection of Microorganisms, culture collection number 4828^T^). *Streptomyces albus* J1074 and *Streptomyces coelicolor* M1152 were used for heterologous expression. All *Streptomyces* strains were grown on soya flour mannitol (SFM) agar medium (20 gL^−1^ soya flour, 20 gL^−1^ mannitol, 20 gL^−1^ agar), with appropriate antibiotic selection upon insertion of plasmid DNA (50 μg mL^−1^ apramycin when transformed with pCRISPomyces2-derived plasmids; 25 μg mL^−1^ kanamycin when transformed with pCAP03-derived plasmids; 25 μg mL^−1^ nalidixic acid on the first round of subculture after intergenic conjugation). *E. coli* ET12567 and ET12567/pUB307 were used for the purpose of intergenic tri-parental conjugation. One Shot TOP10 chemically competent *E. coli* cells (Thermo Fisher Scientific) were used for cloning and storage of plasmid DNA. All *E. coli* strains were grown on lysogeny broth (LB) medium (10 gL^−1^ tryptone, 5 gL^−1^ yeast extract, 10 gL^−1^ NaCl) or LB agar medium (same as LB medium, with 15 gL^−1^ agar), with appropriate antibiotic selection (50 *μ*g mL^−1^ apramycin when transformed with pCRISPomyces2-derived plasmids, 25 *μ*g mL^−1^ chloramphenicol to maintain the *dam* mutation in *E. coli* ET12567, 25 *μ*g mL^−1^ kanamycin either to maintain helper plasmid pUB307 or after insertion of pCAP03-derived plasmids).

*S. cerevisiae* VL6-48N was used for TAR cloning and grown on yeast extract peptone (YPD) broth (5 gL^−1^ yeast extract, 10 gL^−1^ peptone, 2% w/v glucose) or YPD agar (same as YPD, with 15 gL^−1^ agar). Purification of genomic DNA from *S. sclerotialus* was performed from a 100-mL liquid culture by phenol-chloroform extraction.^9^ The *scl* gene cluster was captured using TAR cloning.^23^ Assembly of plasmid pCAP03-scl was performed following the procedure described by Moore and colleagues; pCAP03 was a gift from Bradley Moore (Addgene plasmid # 69862).^24^ For this purpose String DNA fragments (Thermo Fisher Scientific) were ordered to include 60-bp hooks homologous to either side of the *scl* cluster (Supplementary Table S2) and introduced into pCAP03 *via* Gibson Assembly (New England Biolabs). The identity of the captured cluster was confirmed by PCR amplification and restriction digestion (Supplementary Figure S17). Insertion of the *scl* gene cluster in the genome of the heterologous hosts *S. albus* and *S. coelicolor* was accomplished *via* intergenic tri-parental conjugation following the protocol described by Moore and colleagues.^24^ CRISPR/Cas9-based engineering of *S. albus* strains was performed using plasmids pCm2-sclM4, pCm2-sclN, pCm2-sclA and pCm2-sclQ1-4. Golden Gate Assembly was first performed to insert the specific sgRNAs into the backbone pCm2 plasmid, then Gibson Assembly was used to include 800-bp homologous recombination arms, all following the procedure described by Zhao and colleagues (see Supplementary Table S3 for plasmids description); pCRISPomyces-2 was a gift from Huimin Zhao (Addgene plasmid # 61737).^15^ Clearance of temperature sensitive plasmids based on pCm2 was achieved by culturing the mutant strains on SFM agar medium non-selectively at 39°C.

### Identification, isolation and structure elucidation of scleric acid

*S. albus* strains were cultured for 5 days at 30°C on supplemented solid minimal (SM) medium (2 gL^−1^ casaminoacids, 8.68 gL^−1^ TES buffer, 15 gL^−1^ agar; after autoclaving, and just before use, 10 mL of 50 mM NaH_2_PO_4_ + K_2_HPO_4_, 5 mL of 1 M MgSO_4_, 18 mL of 50% w/v glucose and 1 mL of trace element solution [0.1 gL^−1^ each of ZnSO_4_.7H_2_O, FeSO_4_.7H_2_O, MnCl_2_.4H_2_O, CaCl_2_.6H_2_O and NaCl] were added) as described by Hopwood and colleagues.^9^ Ethyl acetate was added in equal volume to the volume of SM medium used, and acidified to pH 3 by the addition of 37% HCl. The ethyl acetate layer was removed and evaporated under reduced pressure. The remaining residue was dissolved in 500 μL of 50:50 (V:V) HPLC grade methanol/water for UHPLC-HRMS analysis. For purification of scleric acid, 2 L of SM culture medium was used for organic extractions, and the remaining residue after evaporation of ethyl acetate was dissolved in 10 mL 50:50 HPLC grade methanol/water for silica column pre-purification.

UHPLC-HRMS analyses were carried out with 20 μL of prepared extracts injected through a reverse phase column (Zorbax Eclipse Plus C18, size 2.1 x 100 mm, particle size 1.8 μm) connected to a Dionex 3000RS UHPLC coupled to Bruker Ultra High Resolution (UHR) Q-TOF MS MaXis II mass spectrometer with an electrospray source. Sodium formate (10 mM) was used for internal calibration and a *m/z* scan range of 50-1500 was used with a gradient elution from 95:5 solvent A/solvent B to 0:100 solvent A/solvent B over 10 minutes. Solvents A and B were water (0.1 % HCOOH) and acetonitrile (0.1 % HCOOH), respectively.

Pre-purification of crude extract containing scleric acid was performed using flash chromatography. A column was loaded with C18-reversed phase silica gel, preconditioned with one volume of methanol, activated with one volume of solvent B (0.045% v/v trifluoroacetic acid in acetonitrile), and equilibrated with two volumes of solvent A (0.045% v/v trifluoroacetic acid in water). Crude extract was loaded onto the column. Compounds were then eluted with five different consecutive solvent systems: two volumes of 20:80 solvent B/ solvent A, two volumes of 40:60 solvent B/ solvent A, two volumes of 50:50 solvent B/ solvent A, two volumes of 60:40 solvent B/ solvent A and two volumes of 80:20 solvent B/ solvent A). Fractions were collected throughout the elution steps, evaporated under reduced pressure and dissolved in 500 μL of 50:50 (V:V) HPLC grade methanol/water for UHPLC-HRMS analysis. Fractions containing scleric acid were combined and used for HPLC purification.

Reverse-phase HPLC was performed using a Zorbax XBD-C18 column (212 × 150 mm, particle size 5 μm) connected to an Agilent 1200 HPLC equipped with a binary pump and DAD detector. Solvent A: 0.1% TFA water, solvent B: 0.1% TFA in acetonitrile, 5% B to 95% B in 45 min. Retention time compound 1: 29.7 min, retention time compound 2: 34.4 min. Gradient elution was used (solvent A: water with 0.1 % HCOOH, solvent B: methanol) with a flow rate of 10 mL min^−1^. Fractions were collected by time or absorbance at 210 nm using an automated fraction collector. The fractions collected containing scleric acid were pooled, methanol removed under reduce pressure and scleric acid was re-extracted from the remaining water (2 × 50 mL ethyl acetate). The ethyl acetate was removed under reduced pressure and the sample re-dissolved in deuterated methanol for NMR analysis.

### MIC testing

The susceptibility of bacterial strains was investigated in collaboration with the Warwick Antimicrobial Screening Facility in a 96-well plate experiment, according to the Clinical & Laboratory Standards Institute (CLSI) guidelines (M7-A9 2012). Scleric acid was diluted to a concentration of 7.5 mg/ml in 25% DMSO in distilled water. To further prevent toxicity effects from DMSO in MIC testing, 27 *μ*1 of the natural product stock was combined with 173 *μ*1 of cation adjusted Muller-Hinton broth to a final concentration of 1024 *μ*g/ml of compound in 200 *μ*1 with 3% DMSO. This was then further doubling diluted throughout the MIC. Meropenem and cefoxitin were used as positive controls during MIC testing.

## Acknowledgements

This project was supported by BBSRC through grant BB/M022765/1, by the Warwick Integrative Synthetic Biology Centre (WISB), a UK Synthetic Biology Research Centre grant from EPSRC and BBSRC (BB/M017982/1) and by The Royal Society via a University Research Fellowship UF090255 to CC. The authors would also like to thank the BBSRC-funded Midlands Integrative Biosciences Training Partnership for a PhD studentship to D. J. Leng. We thank Dr John Sidda and Dr Vincent Poon for preliminary bioinformatics analyses. The Warwick Antimicrobial Screening Facility is thanked. OIDD screening data supplied courtesy of Eli Lilly and Company - used with Lilly’s permission.

## Author contributions

F.A. and C.C. designed experiments and wrote the manuscript. F.A. performed the experiments to capture and activate the cryptic and silent gene cluster. I.W. and L.S. contributed to isolation and characterisation of the novel compound. M.T. designed synthetic routes; D.L. designed and carried out these experiments.

